# Modelling heterogeneities in the cross-linked bacterial sacculus

**DOI:** 10.1101/867176

**Authors:** Garima Rani, Issan Patri

## Abstract

Examining the design principles of biological materials, in particular the presence of inhomogeneities in their ultrastructure is the key to understanding the often remarkable mechanical properties possessed by them. In this work, motivated by the question of understanding the effect of variability in the material properties of the peptide cross-linkers on the bulk mechanical properties of the cell wall structure of bacteria, we study a spring system in which variability is encoded by assigning values of spring constants and rupture strengths of the constituent springs from appropriate probability distribution. Using analytical methods and computer simulations, we study the response of the spring system to shear loading and observe how heterogeneities inherent in the system can heighten the resistance to failure. We derive the force extension relation of the system and explore the effect that the disorder in values of spring constant and rupture strength has on load carrying capacity of the system and failure displacement. We also study a discrete step shear loading of the system, exhibiting a transition from quasi-brittle to brittle response controlled by the step size, providing possible framework to experimentally quantify the disorder in analogous structures. The model studied here will also be useful in general to understand fiber bundles exhibiting disorder in the elasticity and rupture strengths of constituent fibers.

## 1 Introduction

Biological structures are some of the most sophisticatedly engineered materials, whose design principles continue to lend ideas for solving common place as well as esoteric engineering riddles [1, 2, 3]. A pervasive feature of the design of several biological materials is the presence of disorder in their ultrastructural components [4, 5]. However, this cannot be termed as mere accident in light of the often pivotal role played by the disordered fine structure of several key biological materials. Numerous interesting examples of this phenomena can be given, including the role of non-identical molecular motors in actomyosin contractility [6] and the optimized hierarchical structure with highly irregular setup of bone resulting in remarkable orders of toughness and stiffness [7, 4].

One of the most fascinating naturally occurring biomolecule is the peptidoglycan (PG) mesh, the primary component of the cell wall of bacteria. The PG mesh, in rod shaped Gram negative bacteria like *Escherichia coli*, consists of stiff glycan chains arranged roughly in the circumferential direction, cross-linked intermittently by peptide bonds [8, 9] (see Figure 1). It is a testament to the versatility of this structure that it is able to satisfy a wide array of necessary mechanical requirements, including being stiff enough to bear the internal turgor pressure but being elastic enough to allow for elongation, apart from being sufficiently resistant to failure due to crack propagation [10, 8, 9].

**Figure 1:**
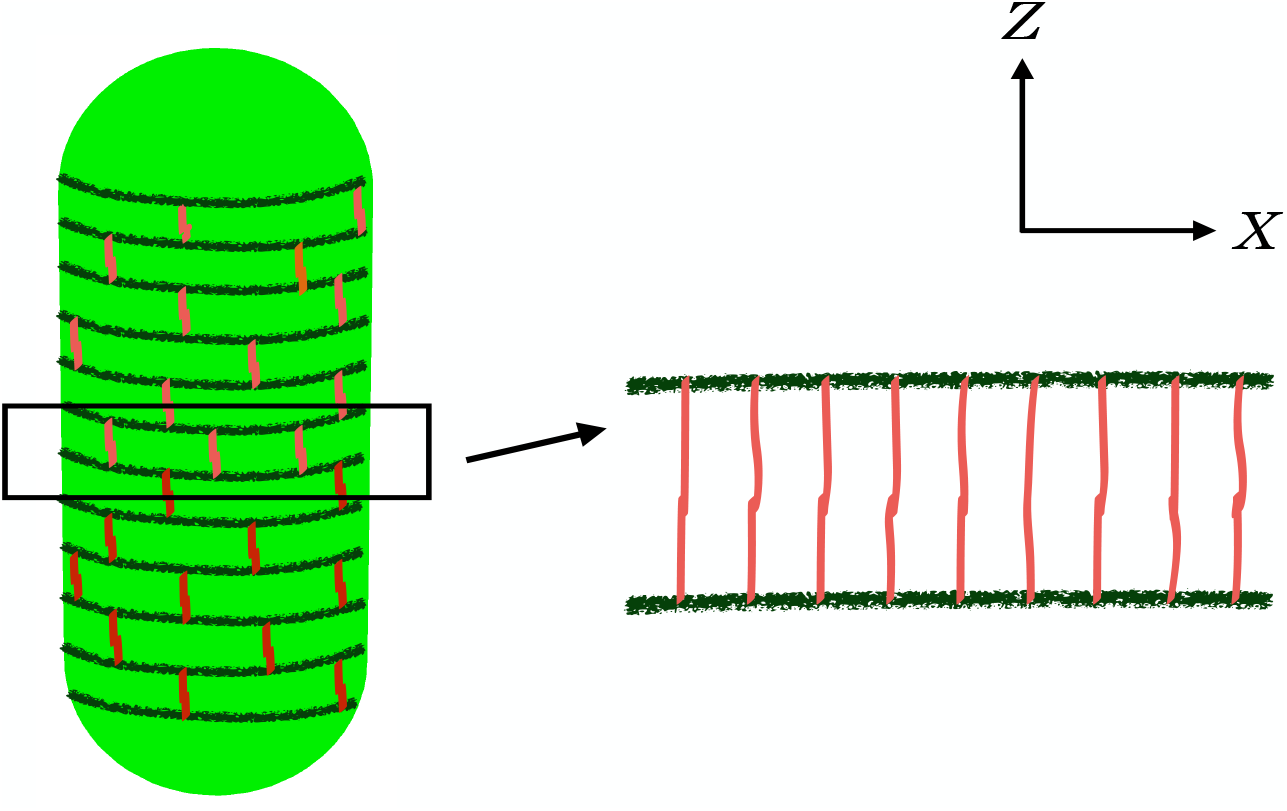
The peptidoglycan (PG) mesh in *E.coli* cell wall has stiff glycan strands aligned roughly in the circumferential direction, crosslinked by short peptide bonds. This mesh can be visualised as an ensemble of stiff interfaces and linear springs.

Like many biological materials, the PG mesh also exhibits disorder that is evident in its ultrastructure- this includes the length distribution and orientation of glycan strands and location and degree of cross-linking across the cell wall [11, 12, 13, 14, 15]. Nonetheless, the relevance of these features of disorder on the mechanical properties of the cell wall has not been well studied, even as it can shed light on a number of experimental observations of the cell wall. For instance, in a previous work [16], we had explored the mechanical effect of length distribution of the glycan strands and had shown how terminally cross-linked smaller length glycan strands can enhance the toughness of the cell wall, giving an explanation of experimental observations of the presence of shorter length glycan strands and the preference for cross-linking to happen at the ends of the glycan strands [11, 14].

The cell wall is continually remodelled in the cell, with cleaving of older cross-links under the action of hydrolases, incorporation of new cell wall material into it and the consequent formation of cross-links [10, 8, 17]. This results in the presence of newer cross-links as well as hydrolase-degraded older cross-links in the cell wall. While the exact mechanical effect of hydrolases on the peptide cross-linkers is unclear, this effect can result in the lowering of rupture strength of the bond or lowering of the stiffness of the bond or both. The presence of heterogeneities in the mechanical properties of the cross-linkers is further evidenced by experimental observations in Ref. [12], where isolated cell wall fragments subjected to sonication showed an immediate drop in the degree of cross-linking that persisted even as the structure remained intact, before becoming relatively constant after a period of time. This suggests that the cross-linkers in the cell wall act in a heterogeneous manner under loading, due to differences in their elasticity and rupture strengths. A very natural question then is to understand effect of the variability in the mechanical properties of the cross-linkers, specifically their strength and elasticity, on the bulk mechanical response of the cell wall. Motivated by this and as a first step, here we explore a spring system akin to a Fiber bundle model (FBM) [18, 19], in which we incorporate variability in both the spring constant of the constituent springs as well as their rupture strengths and study the response of the system to shear loading. We first consider the case when the distribution of the values of the spring constants and rupture strengths are independent. Using analytical methods and computer simulations, we derive the force extension relation and show that in general wider variability in the elasticity of springs can ensure more robust resistance to failure but it also lowers the load capacity whereas lower variability results in a more brittle response to loading even as the load capacity increases. We also deduce that the value of the displacement at failure does not depend on the lower limit of the distribution of the rupture strengths of the springs, though the load capacity of the system increases as the variability in rupture strengths of constituent springs decreases. This suggests that a possible mechanism for hydrolytic action to act on the cross-linkers in a safe manner while ensuring sufficient load bearing ability is by maintaining high order of variability in the spring constants while limiting the variability in the rupture strengths.

Next, we examine a step wise loading regime which allows us to exhibit a quasi-brittle to brittle transition as the load per step increases. This transition, which is a feature that seems universal in systems which have inhomogeneities built into their ultrastructure, has been observed in other natural materials, e.g. snow [20] and can be useful tool to experimentally detect such inhomogeneities present in the system. Finally, we also study the case when the distributions of the spring constants and the rupture strengths of the constituent springs in the system are (positively or negatively) correlated. We show that while in the case of positive correlation, the response to loading is considerably brittle, negative correlation of spring constant and rupture strength values results in quasi-brittle behaviour, although the maximum load carrying capacity drops as compared to independent case. This suggests that to ensure optimal load carrying capacity as well as resistance to failure, it is likely that the spring constant and rupture strength values are independent in natural systems, including the cross-linkers in the peptidoglycan mesh.

## 2 Model

Drawing inspiration from the cross-linked mesh like structure of peptidoglycan in the bacterial cell wall, we study a system of springs placed in the *X* − *Z* plane, consisting of two rigid surfaces, aligned in the *X* direction and linear springs connecting them, aligned in the *Z* direction. The rigid surfaces correspond to two adjacent circumferential cross-sections of the peptidoglycan mesh, consisting of axially adjacent, long length glycan strands, while the springs correspond to the cross-linkers joining the two glycan strands (see Figure 1). The number of springs connecting the upper and lower surface, denoted *N*, is determined by the degree of cross-linking present in the cell-wall-in *E.coli*, observed degree of cross-linking ~ 30% − 50% [10, 14] and with radius of a typical cell ~ 500nm [21], we estimate roughly 1000 peptides stems cross-linked between two adjacent circumferential cross-sections of the cell. Therefore, we take *N* = 1000 in all simulations in this work. Further, since glycan strands are a order of magnitude more stiff than the peptide bonds [22], for simplicity we assume that upper and lower surfaces are rigid with no local deformations caused by exerted forces. The springs in the system are taken to be linear, with finite rupture strength, which limits the force that a spring can endure before rupturing. We encode variability in the system by taking values of the spring constants and the rupture strengths of the springs from an appropriate joint probability distribution over the region [*k*_1_, *k*_2_] × [*f*_1_, *f*_2_] with the upper limits *f*_2_ and *k*_2_, which for illustrative purposes we fix to be equal to 1. Our aim is to tune the range of variability by altering the values of *f*_1_ and *k*_1_ and to see the possible effect on the response of system to loading. In our model, the newly formed cross-links correspond to springs which are stiffer and have higher rupture strengths while older cross-links under effect of hydrolases and mechanical stress, are taken to be relatively weaker and less stiff. Therefore, to tune the range of variability in our model, we vary the values of *f*_1_ and *k*_1_ while keeping *f*_2_ and *k*_2_ fixed. We note that our theoretical setup is independent of the values of the upper limit and our results will not change qualitatively when a change in effected in the values of the upper limit.

In this work, we study shear loading of the spring system. As mentioned before, purified cell wall fragments of *E.coli* have been subjected to sonication [12], a method of cell disruption acting by shear deformation [23] (in general, shearing is a standard and successful method for performing cell disruption experiments [24]). In our case, the upper surface is displaced by application of force while the lower surface is kept fixed (See Figure 2). For simplicity, we assume that adjacent springs maintain the position of their link with the upper and lower surfaces, upto their rupture, ensuring that no sliding motion of the springs occurs. In this case, the elongation is the same for every spring, since the displacement of the point of contact of each spring on the upper surface due to the application of the force, will be the same (equal to displacement of the upper surface). We also take all springs in the system to have the same rest length, equal to the distance between the upper and lower surface (corresponding to the inter-glycan spacing and denoted as *l*, see Fig. 2). In general, though, it is possible that the peptide cross-linkers in the peptidoglycan sacculus have variable rest lengths, which can potentially induce interesting phenomena, for instance emergence of residual stress in the system due to mismatch of the rest lengths of the cross-linkers with inter-glycan spacing. This, and yet other modes of heterogeneity in the peptidoglycan sacculus, will be addressed in a subsequent work.

**Figure 2:**
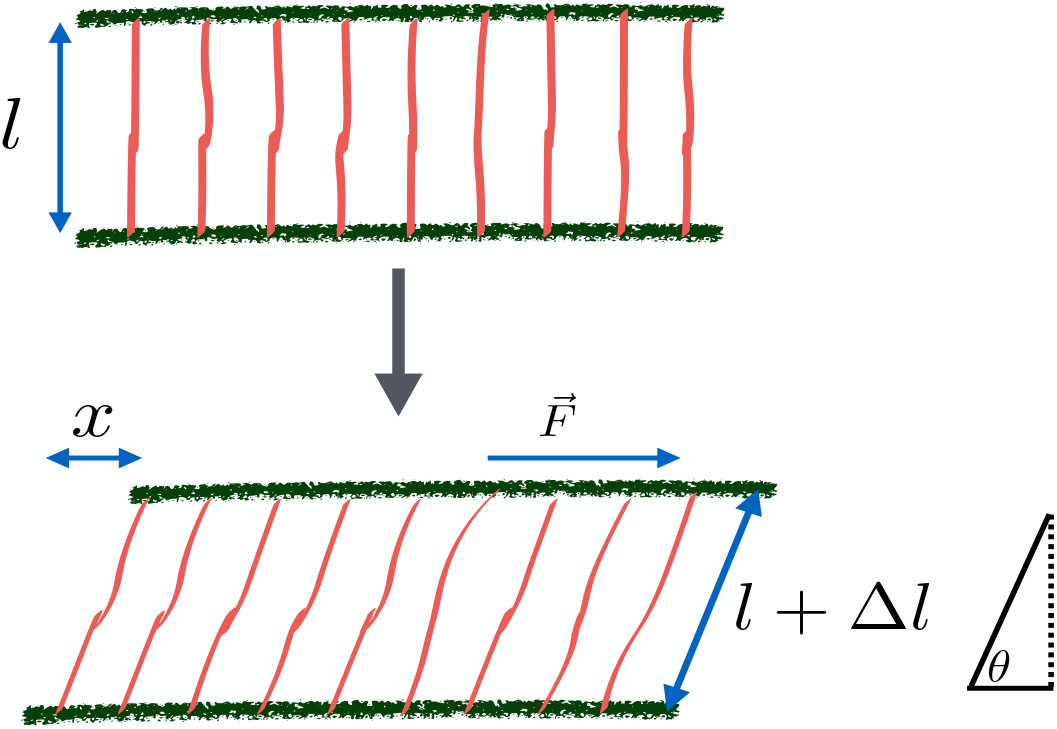
Schematic representation of spring system considered. Two adjacent surfaces at distance *l* from each other joined by *N* linear springs subjected to shear displacement *x*. This results in an extension Δ*l* in each spring, with each spring at an angle *θ* to the lower surface, given by tan *θ* = *l/x*.

### 2.1 Single spring under shear load

We first write down the equations for a single spring with spring constant *k* and rupture strength *f*, under similar shear loading as described above. In general, the system can be loaded in two ways-force controlled loading and displacement controlled loading.

For force controlled loading, we assume that a force *F* is exerted on the upper surface, causing a displacement *x* of the upper surface and resulting in a stretching of the spring by Δ*l*. The force in the direction of spring elongation is then given by *F_θ_* = *k*Δ*l*. Noting that 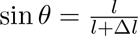 and using force balance, we get

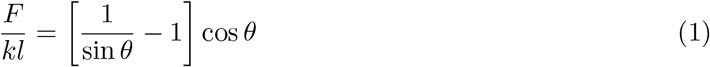

which gives us a polynomial equation for sin *θ* as

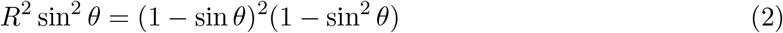

with *R* = *F/*(*kl*). In our case, since 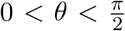, we have 0 < sin *θ* < 1 and it is then easy to see that Eq. 2 has a unique solution. So, solving for sin *θ* using Equation 2, we can calculate the displacement *x* in the *X*-direction as,

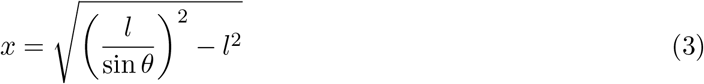

and the shear strain in then given by *ϵ* = *x/l*. In case of displacement controlled loading, a displacement *x* is given to the upper surface keeping the lower surface intact. The extension Δ*l* of the spring is given by

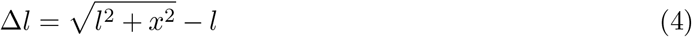

The force *F* inducing the extension *x* on the upper surface is given as

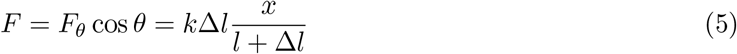

In this work, we carry out a study of displacement controlled loading of the spring system since typically displacement can be measured in an easier and more precise manner.

### 2.2 Constitutive Behaviour of Spring System

We now consider the case of the spring system, undergoing displacement controlled shear loading (see Figure 2), having *N* springs, with spring constants *k*_*i*_, for *i* = 1, 2, ‥, *N*, where N is the total number of peptides between two interfaces. The effective spring constant of the N parallel springs is -

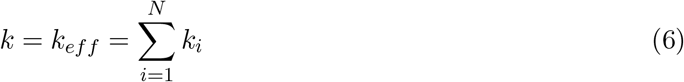

The rupture strength of the springs is denoted *f*_*i*_, *i* = 1, 2, …, *N*. Note that since the shear strain is the same for all (intact) springs, we can calculate the longitudinal elongation of each spring Δ*l* as in Equation 4 and thus the force on the *i*^*th*^ spring is

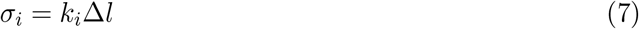

For all springs *i* for which *σ*_*i*_ exceeds the rupture strength *f*_*i*_, the spring breaks. The force being exerted in the *X*-direction on the system is then given by

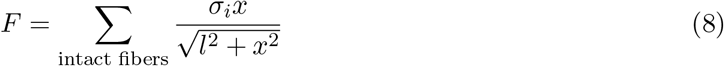

which describes the constitutive behaviour of the bundle.

Consider now the special case where the springs all have same spring constant *k*, the force on all springs is the same, *σ* = *k*Δ*l*, when a displacement *x* is given to the upper surface with Δ*l* as in Equation 4. Further assuming that all the springs have the same rupture strength *f*, the force on the system is given by 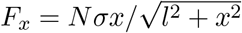, when *σ < f* and the system collapses when *σ* exceeds *f*. However, as mentioned above, it has been observed that cell wall fragments of *E.coli* when subjected to sonication, show an immediate drop in the degree of cross-linking, during which the cell wall structure remains intact [12]. This is however at odds with the behaviour that a system of springs with all springs having same spring constant and rupture strengths exhibits under shear deformation, so we rule out this case and assume that the values of the spring constants and the rupture strengths displays some measure of variability.

To probe the effect of statistical variability in the elasticity and rupture strengths of the springs, we assume that the springs in the system are assigned the values of their respective spring constants and rupture strengths from a joint probability distribution, denoted *p*. So, in particular, the fraction of springs with spring constant in the interval [*a, b*] and threshold values in the interval [*c, d*] is

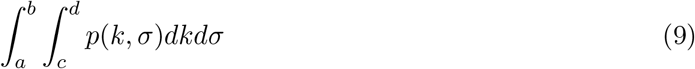

Suppose now that a displacement *x* is imposed on the upper surface keeping the lower surface fixed. The extension in any spring is given by Δ*l*, as given in Equation 4. Since only those springs will survive whose spring constant *k* and rupture strength *f* satisfy *k*Δ*l < f* (see Figure 3), in other words springs whose rupture strength is more than the current load, so the fraction of surviving fibers at given extension *x* is

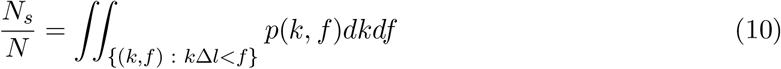

and we have

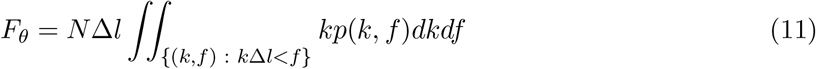

and then the force extension relation is given by *F* = *F_θ_* cos *θ* = *F_θ_x/*(*l* + Δ*l*) as in Equation 5. We note here that in our spring model, the springs, which represent peptide cross-linkers, have been taken to be linear. However, one can study models in which the cross-linkers follow nonlinear force extension relations by suitably modifying our theoretical framework, though this complication is not studied here.

**Figure 3:**
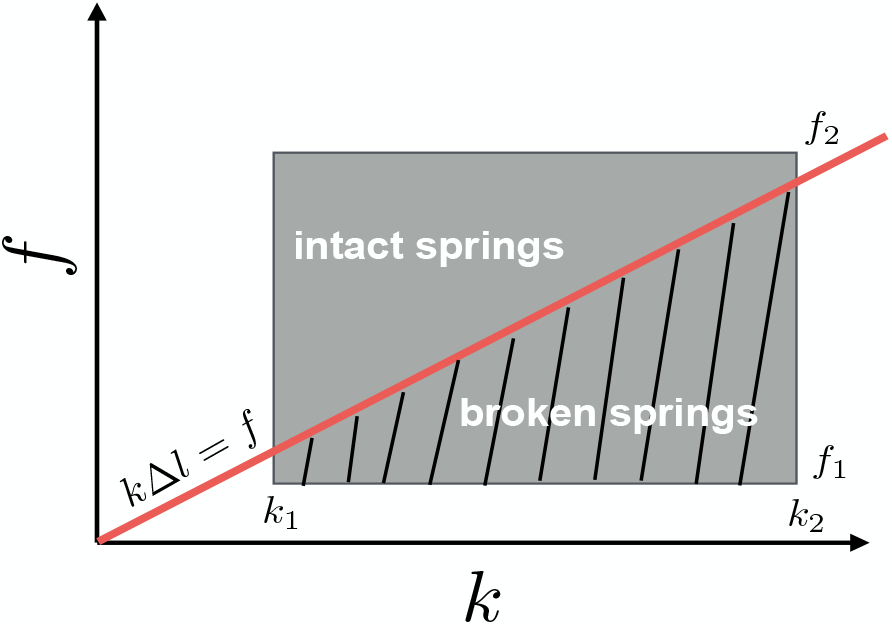
The joint probability distribution of the values of spring constants and rupture strengths of the constituent springs is distributed on [*k*_1_, *k*_2_]×[*f*_1_, *f*_2_]. When a shear displacement *x* is applied to the system, resulting in spring extension Δ*l*, the fraction of broken springs are those whose spring constant and rupture strength lie in the shaded region, below the line *k*Δ*l* = *f*.

### 2.3 Simulation details

We carry out computer simulations to compare with and confirm the theoretical framework laid out in the previous section, for understanding the effect of variability in the material properties of the springs. Specifically, for a given spring system, consisting of *N* springs denoted by *i* = 1, 2, ‥, *N*, the *i*^th^ spring is assigned a tuple (*k*_*i*_, *f*_*i*_), where *k*_*i*_ is its spring constant and *f*_*i*_ denotes its rupture strength. To study the effect of variability on the mechanical properties of the system, values (*k*_*i*_, *f*_*i*_) are drawn randomly from appropriate joint probability distribution for every *i*. Given shear displacement *x*, spring extension Δ*l* is calculated using Equation 4 and the force *σ_i_* on the *i*^th^ spring is then calculated using Equation 7. If *σ*_*i*_ > *f*_*i*_, then the spring is considered ruptured and its spring constant *k*_*i*_ is assigned value 0. The force extension relation is then calculated using Equation 8. The force extension relation and other simulations in the text are obtained by averaging over 100 realizations in each case.

## 3 Results

We now study the effect that the variability in the spring constants and rupture strengths has on the constitutive behaviour of the bundle. We first consider the case where the distributions of the spring constants and rupture strengths of the springs in the system are taken to be independent.

### 3.1 Effect of Variability on Constitutive Behaviour

Since the distributions of the spring constants and rupture strengths are taken to be independent, so the joint probability distribution *p*(*k, f*) (see Equation 9) will decompose as product of the marginal distributions *p*_1_ and *p*_2_, giving *p*(*k, f*) = *p*_1_(*k*)*p*_2_(*f*). We analyse the case where the values of the spring constant and the rupture strength come from uniform distributions. So, with values of the spring constant and the rupture strength lying in [*k*_1_, *k*_2_] and [*f*_1_, *f*_2_] respectively, we get that

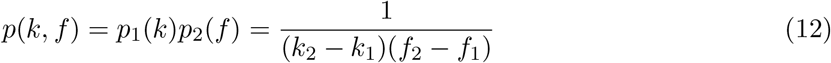

To compute the force extension relation, we note that in this case, the integral in Equation 11 can be evaluated as follows- we define *η* = min(max(*k*_1_Δ*l, f*_1_), *f*_2_). Then we have

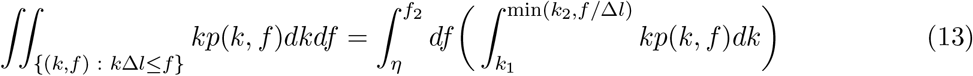

which gives

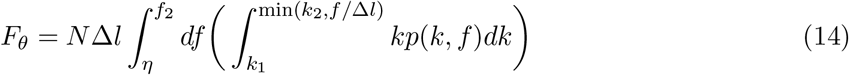

and the force extension relation is then

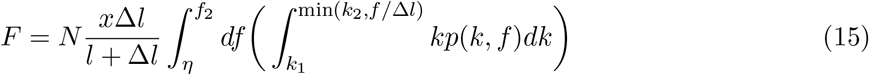

with Δ*l* given as in Equation 4 as a function of the shear displacement *x*.

In Figure 4, using Equation 15 and computer simulations, we compute the force extension curve and the surviving fraction of springs as a function of the shear extension *x*, acting on a fully intact bundle, with the values of spring constants and rupture strengths of the springs in the system drawn from uniform distributions over different intervals. In all cases, simulations show excellent agreement with the theoretical computations. We observe a very interesting contrast-in each case, the maximum load that the bundle takes, given by the maxima of the force-extension curve, is highest in the case when springs in the bundle are the stiffest, given in Figure 4 when the spring constants are uniformly distributed on [0.95, 1]. However, the extension at which the bundle fails, is maximized in the case when the values of the spring constants is distributed over a wider range, which in Figure 4 is given by [0.1, 1], while bundle with stiffest springs show a brittle response to the shear loading. This suggests that while the load bearing ability of the system is enhanced by the presence of stiff springs, the toughness of the system or the resistance of system to mechanical failure is enhanced by the presence of heterogeneities in the system. This also highlights the standard engineering problem of fabricating materials with high degrees of load bearing ability and toughness, something that nature seems to excel in, with biomaterials like bone and nacre exemplifying this property [25, 4]. We also note that the maximum load increases as the variability of the distribution of rupture strength decreases (Figure 4 (c)). Thus, an ideal scenario to ensure high load bearing ability and resistance to mechanical failure is to increase the variation in the value of spring constants while keeping the rupture strength high with little variation. This suggests that a possible mechanism for hydrolytic action on the cross-linkers could result in ensuring wider variability in the elasticity of the cross-linkers while showing little effect on their rupture strengths, so as to secure the viability of the structure which has to bear load even as it is being remodelled continually.

**Figure 4:**
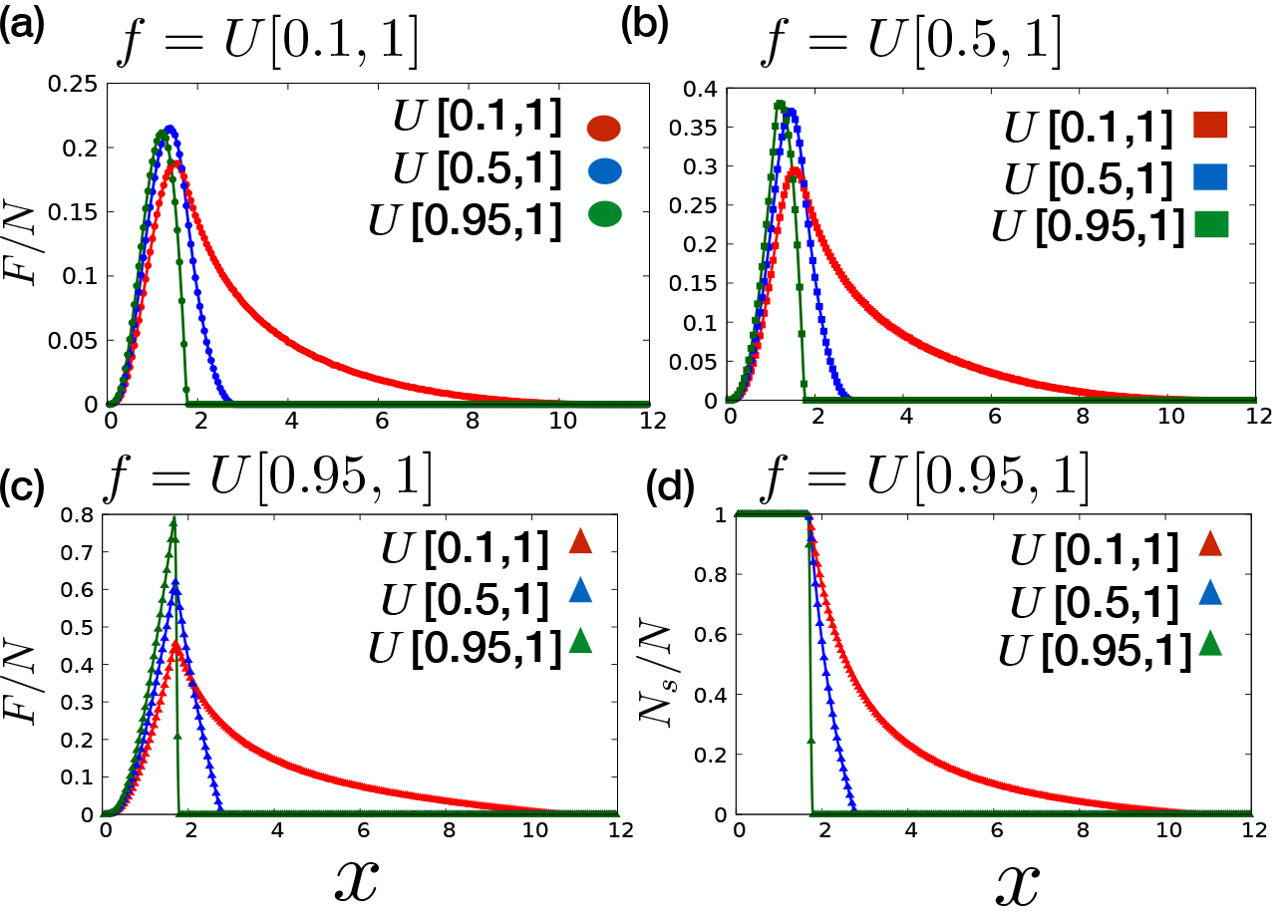
Force-extension curve with distribution of *f* taken in (a) *f* = *U* [0.1, 1], in (b) *f* = *U* [0.5, 1] and in (c) *f* = *U* [0.95, 1]. In each case, three plots are drawn with probability distribution of *k* taken as *U* [0.1, 1], *U* [0.5, 1] and *U* [0.95, 1]. (d) Fraction of surviving fibers is drawn as a function of displacement for *f* = *U* [0.95, 1]. We compare the curve derived analytically from Equations 10 and 15 (solid lines) and simulations (solid points) for all the cases. We have taken *N* = 1000 and *l* = 1.

We now estimate the shear displacement *x*_*i*_ at which spring breakage is initiated and *x*_*f*_ at which the system fails. We observe from Figure 4 (d) that *x*_*i*_ remains unchanged even as the variability in the values of spring constants is changed. This suggests that *x*_*i*_ is independent of *k*_1_. Similarly, we also observe from Figure 4 (a), 4 (b), 4 (c) that *x*_*f*_ stays the same in all three cases once the distribution of spring constant is fixed, which suggests that it is independent of *f*_1_. To understand this interesting phenomena, we estimate *x*_*i*_ and *x*_*f*_ as follows-note that the ratio of rupture strength and the spring constant *f/k* for springs in the system takes value in the range [*f*_1_/*k*_2_, *f*_2_/*k*_1_], since the joint distribution *p* is supported on [*k*_1_, *k*_2_] × [*f*_1_, *f*_2_]. Therefore, breaking of springs is initiated when the displacement ensures that the spring elongation Δ*l* ≈ *f*_1_/*k*_2_ and system failure occurs when Δ*l* ≈ *f*_2_/*k*_1_. We can then estimate the displacement *x*_*i*_ at which spring failure is initiated and the displacement *x*_*f*_ at which the system collapses using Equation 4 which gives

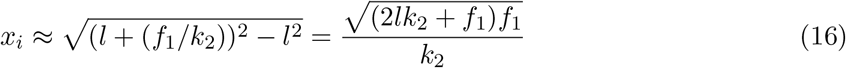

and similarly we have that the bundle fully breaks when Δ*l* = *f*_2_/*k*_1_. So, then we have

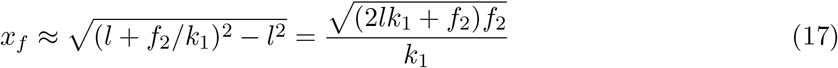

It follows from Equation 17 that *x*_*f*_ depends on the weaker springs that have very high rupture strength, with the extreme case given by springs having spring constant *k*_1_ and rupture strength *f*_2_. It is independent of the values *f*_1_ and *k*_2_, which represent the values for the stiffest springs which have the lowest rupture strength. These values however determine *x*_*i*_ (see Equation 16). Since the displacement is the same for all springs, springs with high stiffness and low rupture strength are the first to rupture. To check this, we perform simulations of bundle of 1000 springs with values of spring constant derived from the distribution *U* [*k*_1_, 1], for varying *k*_1_, while the rupture strength of the springs is derived from fixed uniform distribution, to compute the value of *x*_*i*_ (plotted in Figure 5 (b) as a function of *k*_1_) and *x*_*f*_ (plotted in Figure 5 (d) as a function of *k*_1_). We also perform simulations keeping the distribution of spring constants to be fixed and the rupture strength distribution to be *U* [*f*_1_, 1] with varying *f*_1_ to compute the values of *x*_*i*_ (plotted in Figure 5 (a) as a function of *f*_1_) and *x*_*f*_ (plotted in Figure 5 (c) as a function of *f*_1_). In all cases, the simulations results agrees well with the analytical results. We observe from Figure 5 (b) that the value of *x*_*i*_ is approximately constant in all three cases as *k*_1_ varies, in good agreement with Equation 16, while increasing with value of *f*_1_ approximately linearly. On the other hand, the value of *x*_*f*_ remains constant in all three cases as *f*_1_ varies, as in Figure 5 (c), while showing a power law dependence on *k*_1_. This, in particular, reiterates that the ideal scenario to ensure high value of both *x*_*f*_ and load capacity arises when *k*_1_ has a low value while the value of *f*_1_ is high. This is because a low value of *k*_1_ ensures that *x*_*f*_ has a high value. But the value of *x*_*f*_ remains relatively constant as *f*_1_ changes. Therefore, a high value of *f*_1_, while keeping *x*_*f*_ unchanged, limits the variability and hence, increases the load capacity. To understand this effect, we can make a rough estimation of the shear displacement at failure in case of the bacterial cell wall as the elasticity of the cross-linkers vary-with the spring constant of peptide cross-link estimated as *k*_2_ ~ 10^*−*2^*N/m* [26] and estimating the rupture strength of the cross-link as 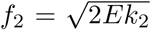*f*_2_ = 2*Ek*_2_ where *E* ~ 300*kJ/mol* is the dissociation energy of the covalent cross-linking bond, we get *x*_*f*_ < 10*nm* when there is very little variation with *k*_1_ ≈ *k*_2_ while *x*_*f*_ ~ 10^2^*nm* when there is variation with *k*_1_/*k*_2_ ≈ 0.1. This calculation, though only a very rough estimation, shows how effectively variation in the elasticity of the cross-linkers can offer high degree of protection from mechanical failure.

**Figure 5:**
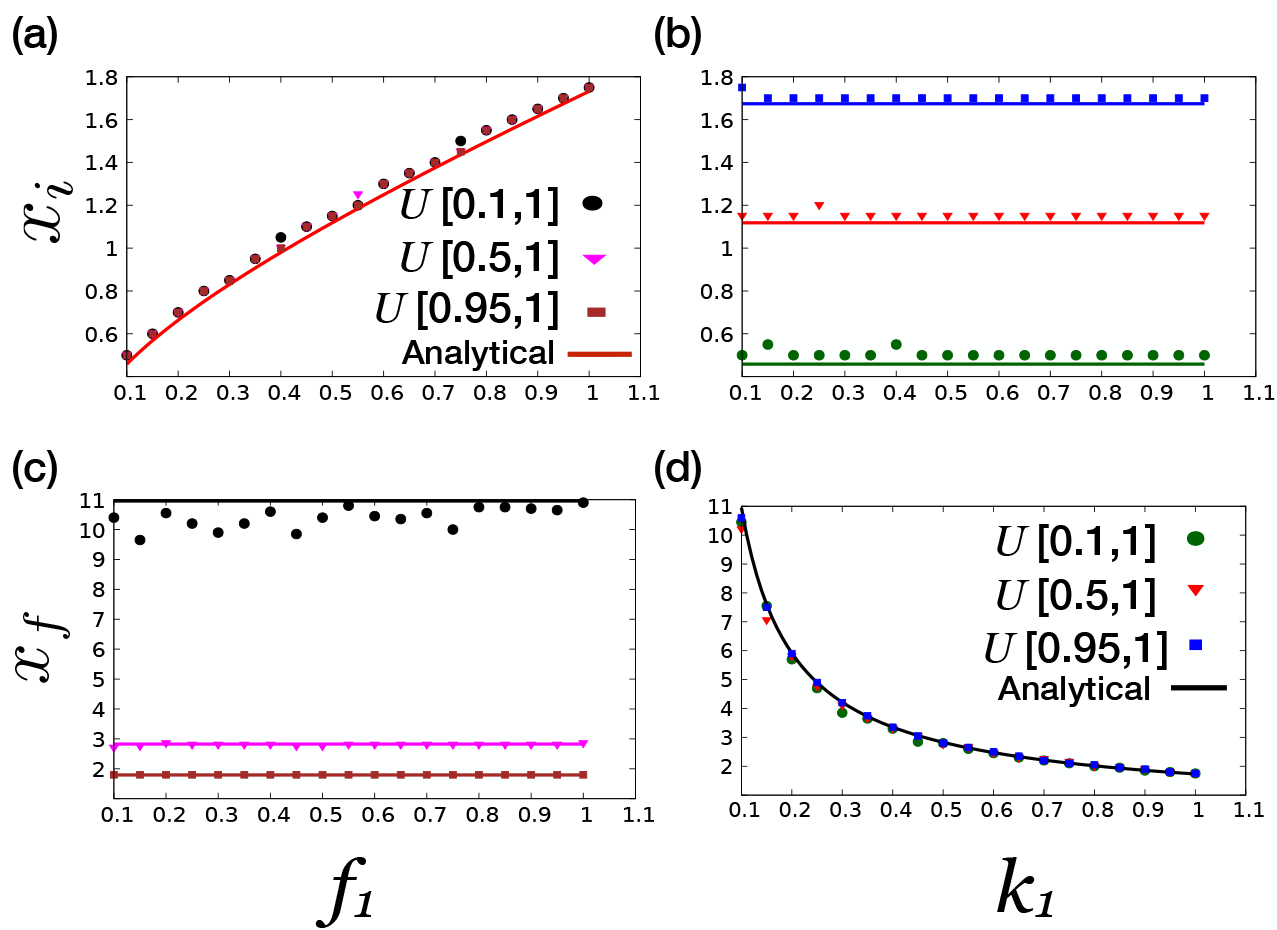
In (a) and (b), displacement at which spring rupturing is initiated, *x*_*i*_, is plotted as a function of *f*_1_, for different distributions of *k*_1_ as labelled and as a function of *k*_1_, for different distributions of *f*_1_, respectively, using Equation 16 (solid line) and simulations (points). In (c) and (d), displacement at which system fails, *x*_*f*_, is plotted as a function of *f*_1_ for different distributions of *k*_1_ and as a function of *k*_1_, for different distributions of *f*_1_ as labelled, respectively, using Equation 17 (solid line) and simulations (points).

Another interesting parameter of the system is the displacement at which the force extension curve attains its maxima, denoted *x*_*m*_. As explained in Appendix A, it is always true that *x*_*i*_ ≤ *x*_*m*_ ≤ *x*_*f*_, it is interesting to observe from Equation 16 and Equation 17 that as the variability in *k* and *f* is limited with *f*_1_ ≈ *f*_2_ and *k*_1_ ≈ *k*_2_, we have that *x*_*i*_ ≈ *x*_*m*_ ≈ *x*_*f*_ in this case, which highlights the decidedly brittle response to loading of the system in this case, with the spring breaking being initiated, the system attaining its load maxima and system failure all occurring when virtually the same shear displacement has been applied to it. As the parameters *f*_1_ and *k*_1_ change, the displacement *x*_*m*_ shows rather intriguing cross-over behaviour- when *f*_1_/*k*_1_ ≪ 1, *x*_*m*_ exhibits independence from the value of *f*_1_ but as *f*_1_ approaches *k*_1_ and becomes greater than it, this behaviour cross-over to one where there is a dependence of *x*_*m*_ on *f*_1_. This behaviour is explored in detail in Appendix A.

### 3.2 Loading Regimes

We now impose specific loading regimes on the spring system and study its response. Specifically, given displacement *x*(*t*) imposed on the upper surface as a function of time *t*, we compute the fraction of surviving springs as a function of time, which we denote *u*(*t*) = *N*_*s*_(*t*)/*N*. We explore two loading regimes-(a) where *x*(*t*) increases linearly with time before becoming constant and (b) where *x*(*t*) increases linearly and then drops to 0 and this cycle is continued, ensuring each time that the load peaks at a level higher than the previous cycle (See Figure 6 (a) and (c)). We note that the response in the first case is an almost immediate drop in the fraction of surviving springs *u*(*t*), which then stabilizes when the loading itself stabilizes. We note the similarity in the response to this loading regime to experimental observations in Ref.[12], where the degree of cross-linking of isolated *E.coli* sacculi subjected to sonication showed an immediate decrease before becoming constant. In the second case, the response to the load results in a calibrated fall in the fraction of surviving springs. When *x*(*t*) is increasing, *u*(*t*) registers a fall but when *x*(*t*) drops to 0 and subsequently increases, *u*(*t*) remains constant till *x*(*t*) goes beyond the previous peak whence *u*(*t*) starts to drop again and this cycle continues.

**Figure 6:**
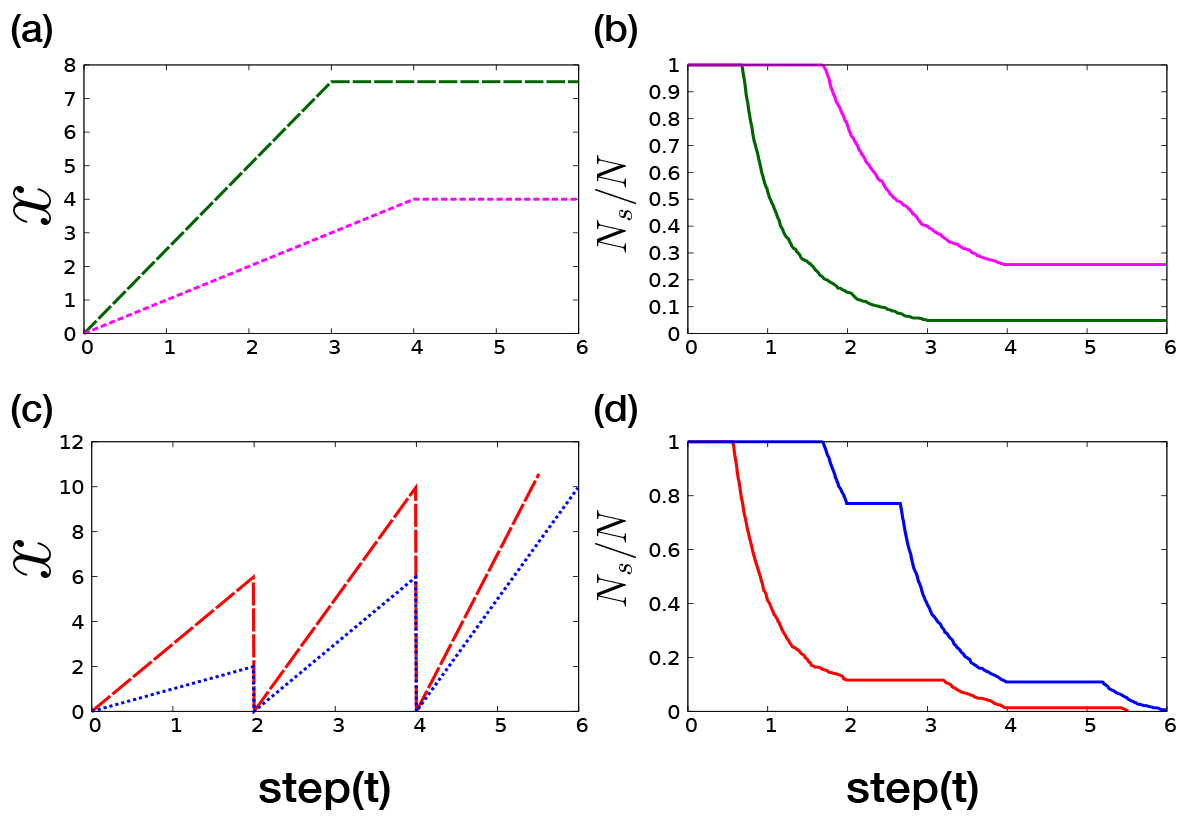
In (a) and (c), loading regimes *x*(*t*) are plotted, given shear displacement at time *t* and the corresponding response of the system to these loading regimes in form of the fraction of surviving fibers is plotted as a function of time in (b) and (d), respectively.

We now consider another loading regime of the spring system, by stepwise increase of the shear displacement. In other words, loading is provided in discrete steps of shear displacement, Δ*x* in each step. We estimate the number of steps to failure in this case. As before, the spring constant and the rupture strength derive values from joint distribution *p*(*k, f*) supported fully on [*k*_1_, *k*_2_] × [*f*_1_, *f*_2_]. The number of steps to failure, denoted *n*_*f*_ (*x*) is given by

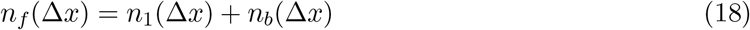

where *n*_1_ denotes the number of steps needed for first failure to happen and *n*_*b*_ denotes the number of steps from first failure till complete failure.

We recall that failure of fibers starts to happen when Δ*l* ≈ *f*_1_/*k*_2_ and all fibers would have failed when Δ*l* ≈ *f*_2_/*k*_1_ (see Figure 3). This gives us the equation

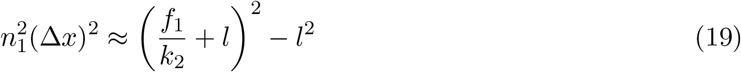

which gives

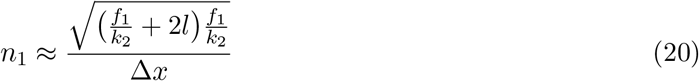

and similarly, we have

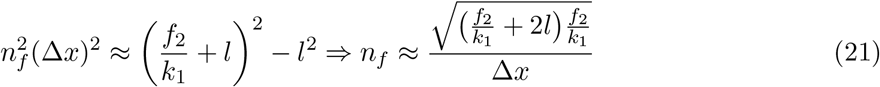

We also define *n*_*b*_(Δ*x*) = *n*_*f*_ (Δ*x*) − *n*_1_(Δ*x*). The response of the system to shear loading is brittle if all springs break in a single step while the response is quasi-brittle in the case when the breaking process occurs over multiple steps. In other words, using notation of Equation 18, the response to shear loading is brittle if *n*_*b*_ < 1 and the response is quasi-brittle if *n*_*b*_ ≥ 1. In Figure 7, we plot the phase diagram of the system in various cases - in Figure 7 (a), the phase diagram is in the *k*_1_-Δ*x* plane while keeping *f*_1_ fixed and in Figure 7 (b), the phase diagram is in the *f*_1_-Δ*x* plane while keeping *k*_1_ fixed. We note that in each case, a quasi-brittle to brittle transition is evident-for sufficiently low values of Δ*x*, the response is quasi-brittle while for high values of Δ*x*, the response is brittle. Now, in case each Δ*x* load is given in fixed Δ*t* time, then the load rate *y* = Δ*x/*Δ*t* can determine the material response-if *y* is high, then the response is brittle as Δ*x* is high as well, similarly, if *y* is low, then Δ*x* is low which results in a quasi-brittle response. This phenomena has been observed to happen in snow [20] and has been studied in [27] using an FBM model that is similar to our model but less general, in that all the springs in the system have a fixed spring constant *k*. In general, this type of quasi-brittle to brittle transition can be considered as a signature of the presence of heterogeneities in the system.

**Figure 7:**
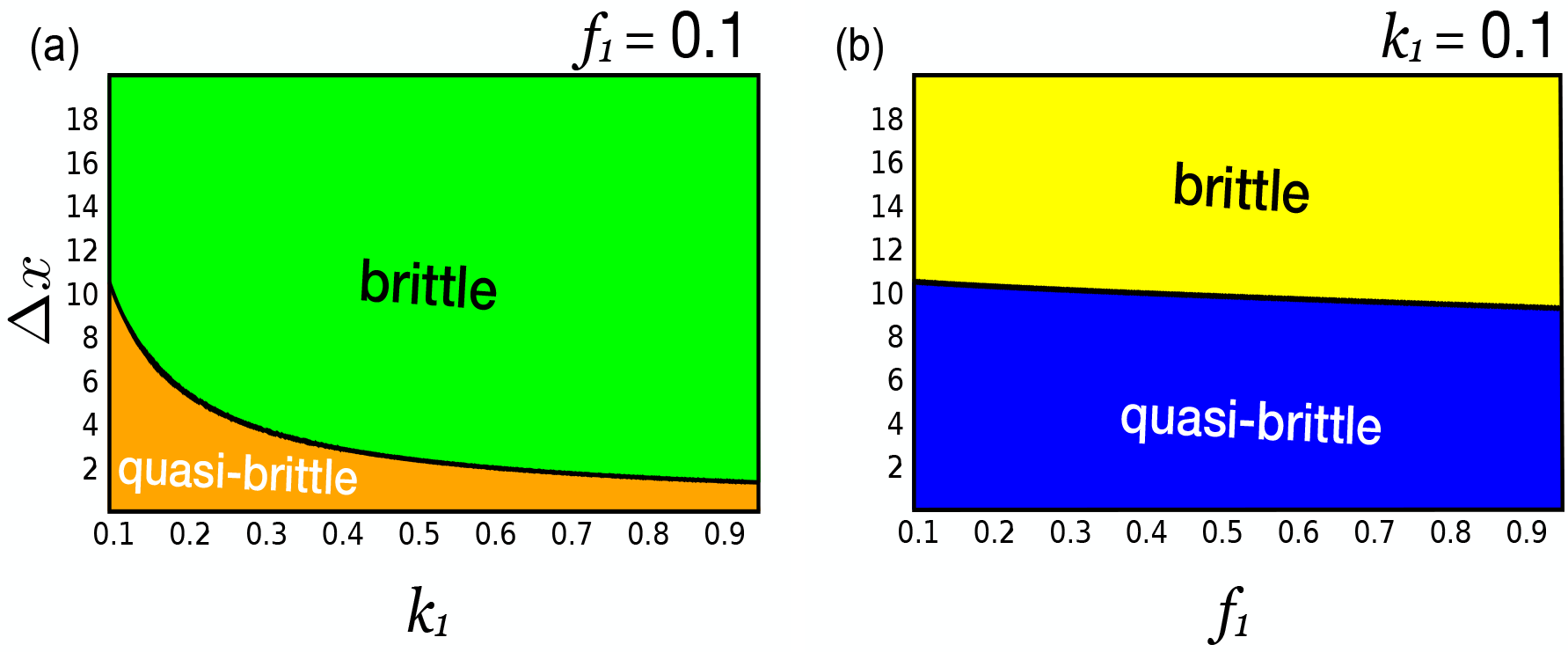
Phase diagram of the spring system with uniformly distributed threshold strength (*f*) and spring constant (*k*), showing a quasi-brittle to brittle transition. (a) Phase diagram in the Δ*x* − *k*_1_ plane for fixed *f*_1_ = 0.1, (b) in Δ*x* − *f*_1_ plane for fixed *k*_1_ = 0.1.

### 3.3 Correlated elasticity and rupture strength

We now study the spring system for which the values of the spring constant and the rupture strength of the constituent springs are correlated. For simplicity, we assume that the values of the spring constant and the rupture strength are derived from the same interval [*a, b*], though we note that our results will hold more generally. We consider first a system with positively correlated *k* and *f* values, with springs having high rupture strength having relatively higher spring constants as well. Fixing *ϵ* > 0, we define the conditional probability distribution of spring constant conditional on the rupture strength 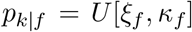, where *U* denotes the uniform distribution on the interval [*ξ_f_, κ_f_*] with *ξ_f_* = max(*a, f* − *ϵ*) and *κ_f_* = min(*f* + *ϵ, b*). Then, with *p*_0_ denoting the probability distribution of the rupture strength and taken as *U* [*f*_1_, *f*_2_], we have the joint distribution as

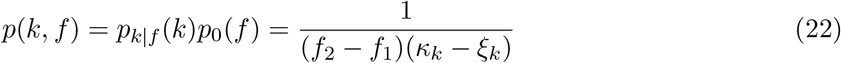

Unlike the previous case, where the spring constants and the rupture strengths of the springs in the system were taken to be independent with the distribution spread fully on the rectangle [*k*_1_, *k*_2_] × [*f*_1_, *f*_2_], in this case, the probability distribution is supported on a strip of width ~ 2*ϵ* around the diagonal (*x, x*), *a* ≤ *x* ≤ *b* of the box [*a, b*]×[*a, b*]. We observe a markedly brittle response to displacement controlled shear loading, as shown in Figure 8, with bundle failure occurring very sharply similar to the case where the spring constants show very little variability while contrasting acutely with the case where the spring constants are spread widely, as in Figure 4 (a).

**Figure 8:**
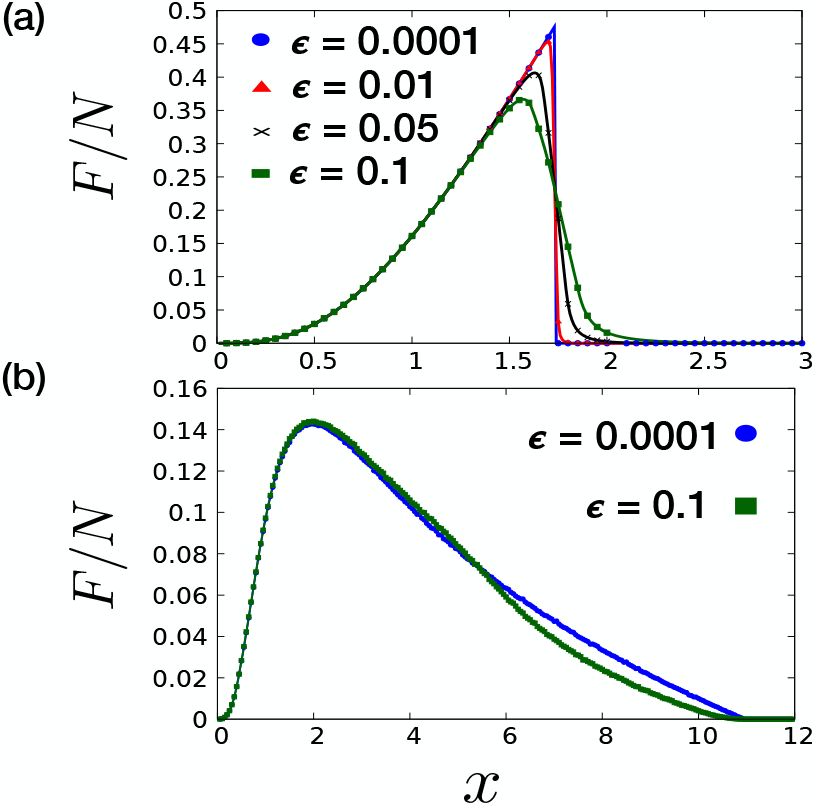
Force extension curve for systems with (a) positively correlated and (b) negatively correlated *k* and *f* values, *ϵ* as labelled in inset, distribution of *f* taken as *U* [0.1, 1] in both cases, computed using Equation 15 (solid lines) and simulations (points).

However, the situation is different when we consider a negative correlation between the *k* and *f* values. We again fix an interval [*a, b*] and *ϵ* > 0. But we now take the distribution *p_k|f_* = *U* [*ρ_f_, μ_f_*], with *ρ_f_* = max(*a, a* + *b* − *f* − *ϵ*) and *μ_f_* = min(*a* + *b* − *f* + *ϵ, b*). We then have the joint distribution *p*(*k, f*) = *p_k|f_* (*k*)*p*_0_(*f*), with *p*_0_ = *U* [*a, b*]. In this case, the values of *k* and *f* are distributed in a region of width ~ 2*ϵ* around the diagonal (*x, a* + *b* − *x*), *a* ≤ *x* ≤ *b*. In Figure 8(b), we observe that the system displays a markedly quasi-brittle response, comparable to the independent case (Figure 4(a)). In other words, the quasi-brittle response to loading of the system with independent *k* and *f* values can be effectively mimicked by systems with *k* and *f* appropriately negatively correlated, even though the support of the values of the spring constants and the rupture strengths of the springs are spread over a much narrower area. To understand this, we note that as the shear displacement is increased, the region [*a, b*] × [*a, b*] in the *k* − *f* plane is swept in the anti-clockwise direction by lines of the form *f* = *k*Δ*l*, with the area under the line determining the fraction of springs broken (see Figure 3). In case the values of *k* and *f* are negatively correlated, the entire area of [*a, b*] × [*a, b*] has to be swept to cover all the springs. In the positively correlated case, the *k* and *f* values are distributed around the diagonal (*x, x*), so much lesser area has to swept to cover all springs in the system, hence it exhibits a decidedly brittle response. However, while in the positive correlation case the maximum load is greater than the independent case, in case of negative correlation, the maximum load is in fact lower than the independent case. Hence, our analysis suggests that, in general, for ensuring a sufficiently high load capacity and also high resistance to failure, it can be important that the distributions of *k* and *f* of the springs in the system are independent and are supported over a wide region, essentially resulting in a multi-composite structure. For the particular case of the bacterial cell wall, it is therefore plausible that hydrolytic action effects independent distributions of the elastic properties of the peptide cross-linkers, with wide variability in the stiffness and limited variability in rupture strengths, since this has the effect of making the structure both more resistant to failure under to loading and enhances the load carrying capacity.

## 4 Non uniform distributions

So far, we have primarily considered the case with spring constants and rupture strengths of the springs in the system having uniform distribution over appropriate intervals. We now study systems for which the *k* and *f* values are independent and follow other distributions. First, we consider the case where the spring constants and the rupture strengths follow Gaussian distributions *N* (*μ*_0_,*σ*_0_) and *N* (*μ*_1_, *σ*_1_) respectively. For a random variable following the *N* (*μ, σ*) distribution with mean *μ* and standard deviation *σ*, it is easy to see that ≈ 99.7% of values lie within a distance of 3*σ* of the mean *μ*. We therefore infer that the spring constants and rupture strengths of the constituent springs in the system are essentially distributed in the region [*k*_1_, *k*_2_] × [*f*_1_, *f*_2_] with *k*_1_ = *μ*_0_ − 3*σ*_0_, *k*_2_ = *μ*_0_ + 3*σ*_0_, *f*_1_ = *μ*_1_ − 3*σ*_1_, *f*_2_ = *μ*_1_ + 3*σ*_1_. Given *k*_1_, *k*_2_, *f*_1_, *f*_2_, we solve and get *μ*_0_ = (*k*_1_ + *k*_2_)/2, *σ*_0_ = (*k*_2_ − *k*_1_)/6, *μ*_1_ = (*f*_1_ + *f*_2_)/2, *σ*_1_ = (*f*_2_ − *f*_1_)/6. So, we have

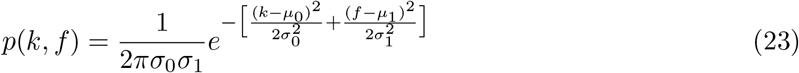

The force extension relation is then computed using Equation 15 (noting that the integral in this case cannot in general be exactly solved, unlike the uniform case and is therefore numerically approximated) and it is compared with computer simulations of the system as described in Section 2.3, in Figure 9. Simulations show excellent agreement with theoretical computations, which demonstrates that our theoretical framework is applicable for spring systems irrespective of the distributions followed by *k* and *f* values.

**Figure 9:**
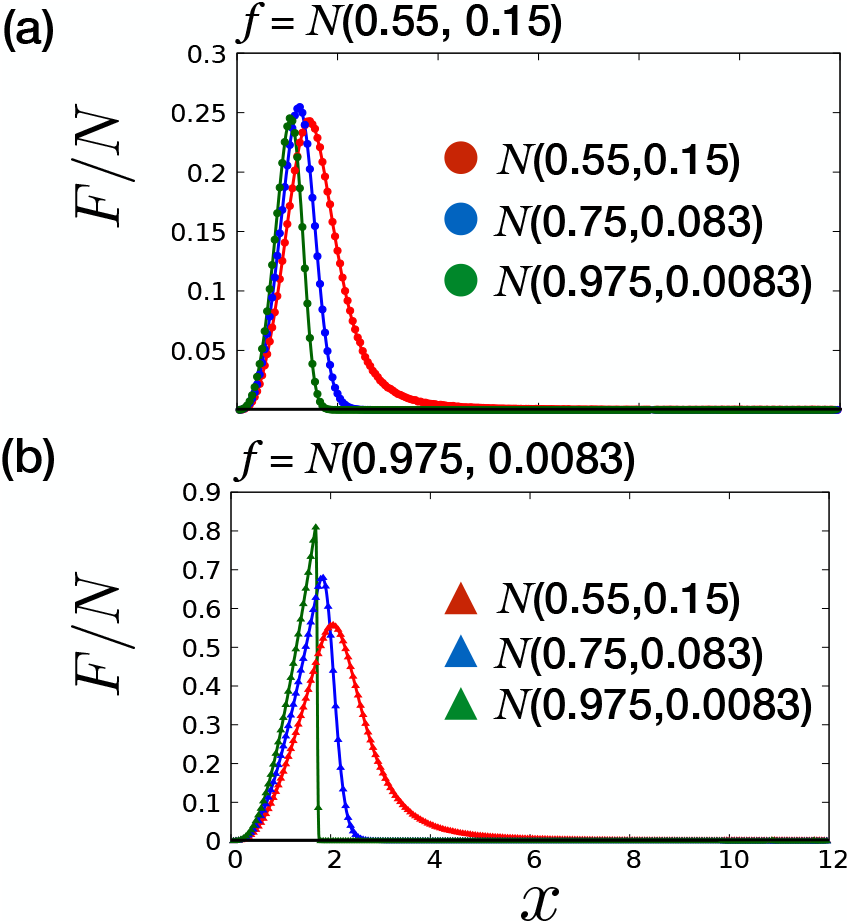
Force extension curve for systems with *k* and *f* values following independent Gaussian distributions. The *f* values follow (a) *N* (0.55, 0.15) and (b) *N* (0.975, 0.0083) distributions while the distributions of *k* values are labelled in inset. We compare the curves derived using Equation 15 (solid lines) and simulations (points).

In this case, we note that 1-the maximum load is always higher in the Gaussian case as compared to the uniform case and 2-the system in this case collapses at a faster rate, with a significant number of the springs having ruptured at shear displacements much lower than *x*_*f*_. This is because in the Gaussian case the *k* and *f* values are strongly concentrated around the mean but in the uniform case the values are spread more evenly. To further highlight this, we also consider systems for which the spring constants follow a left truncated and a right truncated normal distributions, essentially supported in [*k*_1_, *k*_2_], with probability distributions given by

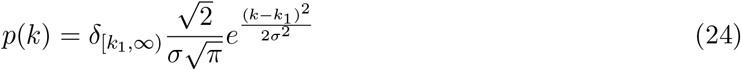

and

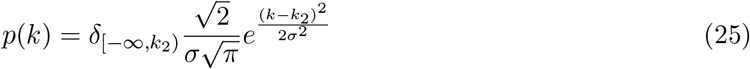

respectively (see Fig. 10 (inset)). Here *σ* = (*k*_2_ − *k*_1_)/3 and the function *δ*_[*a,b*]_(*x*) takes value 1 for *a* ≤ *x* ≤ *b* and 0 otherwise. In both the uniform and Gaussian cases, wider variability in the *k* values and limited variability in the *f* values results results in an ideal scenario with a high load carrying capacity and resistance to failure, so we fix this case with *f* values showing relatively much less variability and following the uniform distribution on a small interval. In Figure 10, we compare the force extension curves with the spring constants having uniform, Gaussian, left truncated Gaussian and right truncated Gaussian distributions. We observe that the maximum load is highest for the right truncated case and is lowest in the left truncated case, implying that the load capacity is higher when the spring constants are concentrated around a higher value. However, while the systems finally fail at same shear displacement *x*_*f*_ for all distributions, the right truncated system fails at the fastest rate and the left truncated system displays a force extension curve that is the slowest to fall, exhibiting substantial quasi-brittle response to loading.

**Figure 10:**
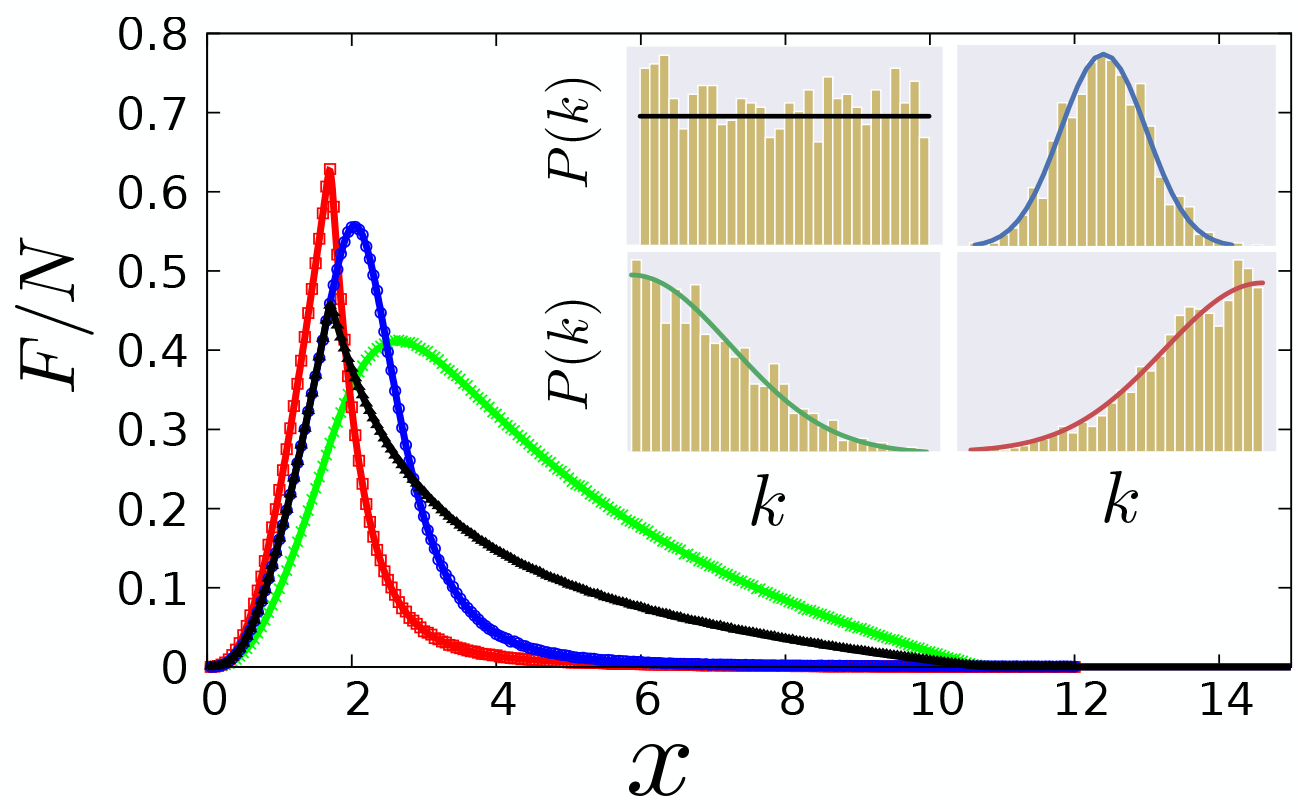
Force extension curves for systems with *k* and *f* values taken independent, with distribution of *f* fixed as *U* [0.95, 1], computed using Equation 15 (solid lines) and simulations (points). The *k* values follow the uniform (black), Gaussian (blue), left truncated Gaussian (green), right truncated Gaussian (red) distributions, supported on [0.1, 1]. The distributions are depicted in the inset.

We note here that the main results and conclusions of this study are not dependent on the form of the distributions of the values of *k* and *f*, however, the precise shape of the force extension curve will depend on the distributions. In general, the distributions and the range of the *k* and *f* values in the peptidoglycan mesh will be influenced by the rate of incorporation of cell wall material (synthesis rate) and the rate at which hydrolases act to facilitate material removal in the cell wall, factors which modulate the growth rate of the cell as well [28, 29]. For instance, uniform distribution distribution of the values of *k* and *f* will likely occur under conditions resulting in steady incorporation and removal of cell wall material. On the other hand, a concentration of the *k* and *f* values closer to the upper limit of the distribution is probably indicative of a higher rate of synthesis as compared to the rate of hydrolysis while a faster rate of hydrolysis might result in a distribution of values concentrated near the lower limit. Since dissimilar growth conditions will likely result in elastic properties of the peptide cross-linkers being differently distributed, our analysis thus suggests that the cell wall adopts disparate constitutive behaviour under varying growth conditions. A detailed theoretical study of this will be done in a subsequent work. Another interesting direction in this regard is the possibility of carrying out a form of biological “forensics”, to ascertain by carefully analysing constitutive behaviour of isolated cell wall fragments, the conditions undergone by the cell itself during its growth.

## 5 Discussion and Conclusion

In this work, inspired by the cross-linked structure of the bacterial cell wall and to understand the effect of variability in the mechanical properties of the peptide cross-linkers on the structure of the cell wall, we studied the response of a spring system, consisting of several springs joining two adjacent rigid surfaces, to a displacement controlled shear loading. Variability in the mechanical properties of the cross-linkers has been indicated by experimental results on sonication of isolated *E.coli* sacculi [12] and can arise from action of hydrolases on the cell wall, resulting in a distribution of newly formed and degraded cross-links. To incorporate variability into the model, the spring constants and rupture strengths of the springs were taken from an appropriate probability distribution. Laying the condition that the distribution of spring constants and rupture strengths are independent, we computed the force extension curve and observed that higher variability in values of the spring constants resulted in a quasi-brittle response to loading with a higher value of the shear displacement at which the system collapses while lower variability resulted in a much more brittle response, highlighting the standard problem in engineering of ensuring high orders of stiffness and toughness in a material [1]. On the other hand, while the load capacity increased with lower variability in values of rupture strength, the failure displacement remained independent of the lower limit of its distribution. Thus our work reflected a viable way for providing a high load bearing capacity and resistance to failure to the system is by ensuring a composite structure with wide variability in value of spring constants and low variability in rupture strengths. Since the bacterial cell wall is key to bearing turgor pressure in the cell while being remodelled, with cleaving of cross-links and insertion of newer glycan strands happening continually, maintaining structural integrity under such conditions is critical to the survival of the cell. Our work suggests that the action of hydrolases, if resulting in good variability in the elasticity of cross-linkers while showing little effect on their rupture strength, can ensure that the structure remains robust enough to sustain high turgor pressure as well as resist mechanical failure.

We also explored a stepwise loading regime which allowed us to obtain a quasi-brittle to brittle transition as the load increases. This transition is a signature of the presence of heterogeneities in the system and hence, is of much importance for experimental detection of the same.

In this work, we have modelled the peptide cross-linkers in the peptidoglycan sacculus as linear springs. Although peptide cross-linkers have often been modelled as linear springs in previous work [26, 29, 30], non-linearity in their force extension relation is plausible. Indeed, simulation work has suggested that their force extension curves are well approximated as a worm like chain (WLC) [31]. We note that our theoretical framework can be suitably modified to incorporate non-linearity in the force extension relation of the peptide cross-linkers, nevertheless, the main results of our study will not change qualitatively.

Several theoretical approaches to modelling the cell wall have taken a continuum theory approach [32, 33, 34], which however may not take into account the molecular level architecture and the inhomogeneities inherent in the structure of the cell wall. The peptidoglycan sacculus has a significantly complex structure and we acknowledge that in our simplified model, we have not considered and studied the full range of its design features and their role in ensuring the stability of the cell wall and survival of the cell. Nonetheless, our approach presents a possible first step towards including the inhomogeneities present in the peptidoglycan mesh and its molecular scale architecture in modelling of the cell wall structure and dynamics. In particular, our work anticipates a potentially paradigmatic shift in the coarse grained modelling of the peptidoglycan sacculus by considering variability in the elastic properties of the constituents, leading to a better understanding of the bulk material properties of the sacculus - previous coarse grained models of peptidoglycan have typically assumed uniformity in the elastic properties of glycan strands and peptide cross-linkers [26, 29, 30]. Future work will involve incorporating the dynamics of the cell wall and the elasticity of the glycan strands into our model alongwith local transfer of load when a cross-link ruptures, from which stress concentrations and pore size distributions can be computed and compared with experiments and simulations, which can lead to further insights on the structure of the cell wall and shed light on its surprising viability against all odds.

**Figure 11:**
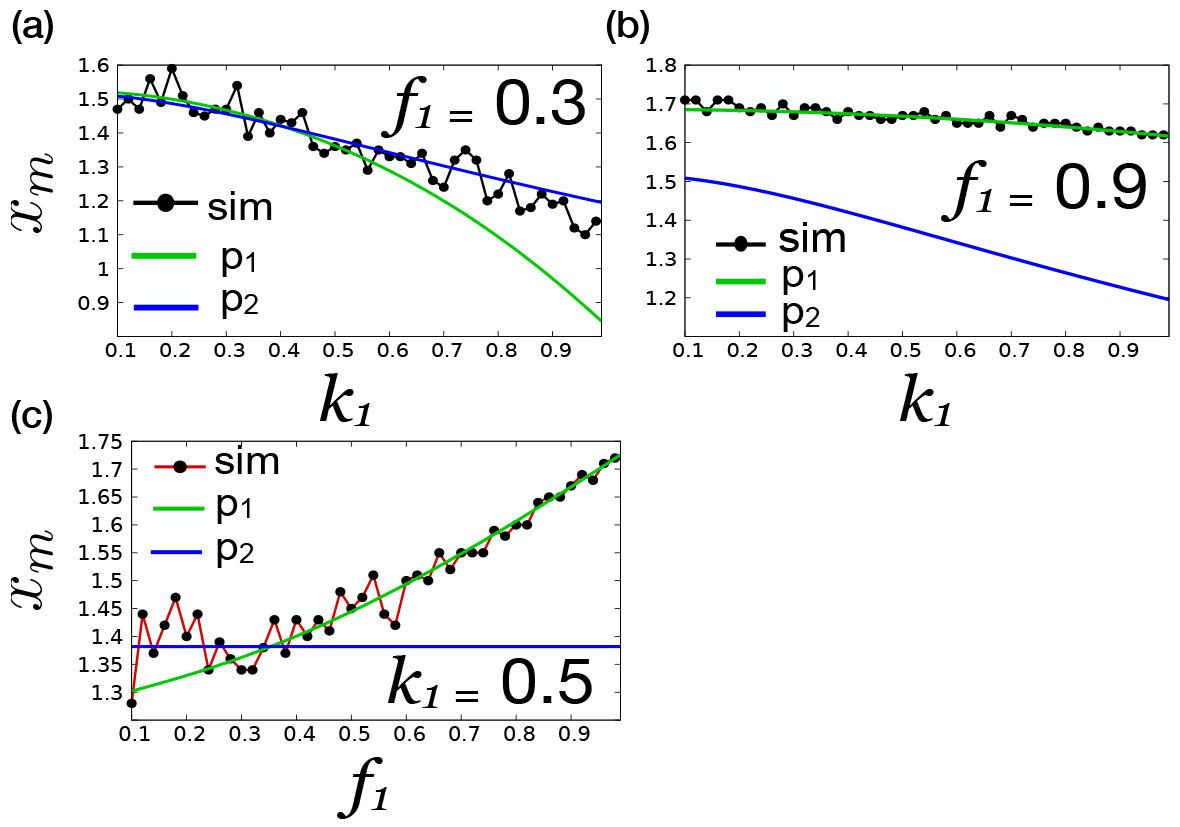
In (a) and (b), the displacement for which the force extension curve attains maxima, *x*_*m*_, is plotted as a function of *k*_1_, with fixed *f*_1_ = 0.3 and fixed *f*_1_ = 0.9 respectively. In (c), *x*_*m*_ is plotted as a function of *f*_1_ with fixed *k*_1_ = 0.5. The green and blue solid line indicate the value of *x*_*m*_ derived from Δ*l* taken as the unique solution of polynomials *p*_1_ and *p*_2_ while dotted line indicates *x*_*m*_ obtained from simulations.

## A Shear Displacement and Maximum Load Capacity

Here we analyse the shear displacement *x*_*m*_ at which the force extension curve attains its maxima as a function of the parameters *f*_1_ and *k*_1_, with *f*_2_ = *k*_2_ = 1. Since the shear displacement uniquely determines the spring extension Δ*l* by Equation 4 in the main text, the force extension relation here is taken as a function of Δ*l*. There are two possibilities-(a) *f*_1_/*k*_1_ ≤ *f*_2_/*k*_2_ and (b) *f*_2_/*k*_2_ ≤ *f*_1_/*k*_1_. In case (a), it follows from a straightforward calculation using Equation 15 in the main text, that the derivative of *F* with respect to Δ*l*, *F′* (Δ*l*) > 0 when Δ*l < f*_1_/*k*_2_ and that *F′*(Δ*l*) < 0 when *f*_2_/*k*_2_ = 1 < Δ*l < f*_2_/*k*_1_, with *F* ≡ 0 when Δ*l* ≥ *f*_2_/*k*_1_. In other words, we have that *F* strictly increases as a function Δ*l* when Δ*l* increases between 0 and *f*_1_/*k*_2_ while it strictly decreases when Δ*l* increases between *f*_2_/*k*_2_ = 1 to *f*_2_/*k*_1_ beyond which *F* ≡ 0. Therefore, in this case, maxima is attained either when *f*_1_/*k*_2_ ≤ Δ*l* ≤ *f*_1_/*k*_1_ or when *f*_1_/*k*_1_ ≤ Δ*l* ≤ *f*_2_/*k*_2_ = 1. With *f*_1_/*k*_2_ ≤ Δ*l* ≤ *f*_1_/*k*_1_, the force extension relation as in Equation 15 in the main text, becomes

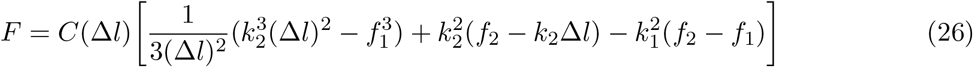

where we have

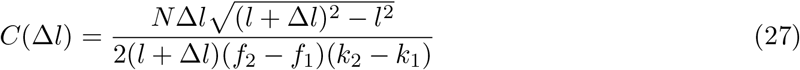

Now, to compute *x*_*m*_, we differentiate *F* as given by Equation 26 with respect to Δ*l* and equating with 0, we get the fifth order polynomial equation, denoted *p*_1_(Δ*l*) = 0, given by

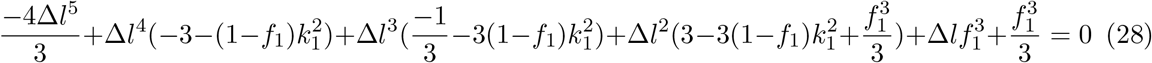

Next, when *f*_1_/*k*_1_ ≤ Δ*l* ≤ *f*_2_/*k*_2_ = 1, we have

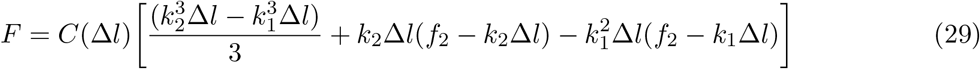

with *C*(Δ*l*) as in Equation 27. Again, we differentiate *F* with respect to Δ*l* and equating with 0, we get a cubic equation, denoted *p*_2_(Δ*l*) = 0, given by

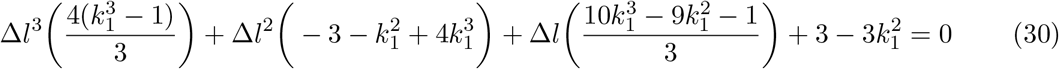

Interestingly, the cubic polynomial *p*_2_ is independent of *f*_1_ and thus, in case that the maxima is attained when *f*_1_/*k*_1_ ≤ Δ*l* ≤ 1, then the shear displacement at which maxima is attained is independent of *f*_1_. Since *p*_1_(0) > 0 and *p*_2_(0) > 0 while *p*_1_(1) < 0 and *p*_2_(1) < 0, this implies that both *p*_1_ and *p*_2_ have roots in the interval [0, 1]. It follows from discriminant analysis and by numerical methods that both *p*_1_ and *p*_2_ have unique root in this interval. So, the question arises as to when the root of *p*_1_ lies in the interval [*f*_1_/*k*_2_, *f*_1_/*k*_1_] and likewise, when the root of *p*_2_ lies in the interval [*f*_1_/*k*_1_, 1]. We note that

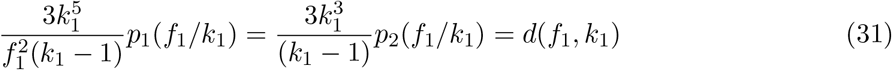

where *d*(*f*_1_, *k*_1_) is the following expression in *f*_1_ and *k*_1_-

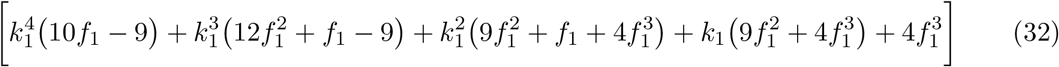

Since *p*_1_(*f*_1_/*k*_2_) > 0, *p*_2_(1) < 0 and *k*_1_ < 1, it follows that if *d*(*f*_1_, *k*_1_) > 0 then *p*_1_ has a root in the interval [*f*_1_/*k*_2_, *f*_1_/*k*_1_] and consequently the maxima of the force extension curve lies in this interval. On the other hand, if *d*(*f*_1_, *k*_1_) < 0, then *p*_2_ has a root in the interval [*f*_1_/*k*_1_, 1], therefore the force extension curve attains maxima in this interval, in which case, as mentioned, *x*_*m*_ is independent of the value of *f*_1_ since *p*_2_ is independent of *f*_1_. We note that when *k*_1_ ≈ *f*_1_, then *d*(*f*_1_, *k*_1_) > 0, which implies that in this case, *p*_1_ has a root between [*f*_1_/*k*_2_, *f*_1_/*k*_1_]. Similarly, for high values of *f*_1_, for instance, with *f*_1_ ≥ 0.8, we also have *d*(*f*_1_, *k*_1_) > 0, in which case *x*_*m*_ is derived from the root of *p*_1_ in interval [*f*_1_/*k*_2_, *f*_1_/*k*_1_]. But when *f*_1_/*k*_1_ ≪ 1, then *d*(*f*_1_, *k*_1_) < 0 and so, *x*_*m*_ is derived from the root of *p*_2_ in the interval [*f*_1_/*k*_1_, 1].

Next, in case when 1 = *f*_2_/*k*_2_ ≤ *f*_1_/*k*_1_, as before we have *F′*(Δ*l*) > 0 for any 0 < Δ*l < f*_1_/*k*_2_ and *F′*(Δ*l*) < 0 when *f*_2_/*k*_2_ = 1 < Δ*l < f*_2_/*k*_1_ with *F* ≡ 0 for Δ*l* ≥ *f*_2_/*k*_1_. Therefore, maxima is attained when *f*_1_/*k*_2_ ≤ Δ*l* ≤ *f*_2_/*k*_2_ = 1. In this case, Δ*l* is also a root of *p*_1_ as given in Equation 28 and is the unique root in interval [*f*_1_/*k*_2_, 1]. In Figure 11, we plot *x*_*m*_ as a function of *k*_1_ fixing *f*_1_ and of *f*_1_ fixing *k*_1_. Specifically, in Figure 11 (a) with fixed value of *f*_1_ = 0.3, when *k*_1_ ≤ *f*_1_, we have that 1 = *f*_2_/*k*_2_ ≤ *f*_1_/*k*_1_ and so, *x*_*m*_ is derived from the root of *p*_1_ (shown in green color). However, when *k*_1_ ≥ *f*_1_, except for a crossover region where *f*_1_ ≈ *k*_1_ in which the shear displacement at maxima is derived as a root of *p*_1_, for larger values of *k*_1_ with *f*_1_/*k*_1_ ≪ 1, we see that *x*_*m*_ is derived from the root of *p*_2_ (blue color in Figure 11). In Figure 11 (b), a high value of *f*_1_ = 0.9 is fixed and as discussed, in this case, *x*_*m*_ is derived from the root of *p*_1_ for all values of *k*_1_. In Figure 11 (c), we have fixed value of *k*_1_ = 0.5. We note that for smaller values of *f*_1_, with *f*_1_/*k*_1_ << 1, *x*_*m*_ is derived from the root of *p*_2_ and is therefore, constant as *f*_1_ varies. However, as *f*_1_ approaches *k*_1_, the behaviour crosses over with *x*_*m*_ now derived from the root of *p*_1_, and as *f*_1_ becomes greater than *k*_1_, we have *f*_1_/*k*_1_ > 1 and as discussed, the *x*_*m*_ is still derived from the root of *p*_1_. In all cases, simulation results show excellent agreement with theoretical results.

## Acknowledgements

The authors thank B.V. Rao and Himalay Senapati for useful suggestions. I.P. is supported by a DST Inspire Faculty Award (DST/INSPIRE/04/2016/001185).

